# AFM characterization of the interaction of PriA helicase with stalled DNA replication forks

**DOI:** 10.1101/2020.02.12.945972

**Authors:** Yaqing Wang, Zhiqiang Sun, Piero R. Bianco, Yuri L. Lyubchenko

**Author notes:** To whom correspondence should be addressed: Yuri L. Lyubchenko: Department of Pharmaceutical Sciences, College of Pharmacy, University of Nebraska Medical Center, 986025 Nebraska Medical Center, Omaha, NE 68198-6025.

## Abstract

In bacteria, the restart of stalled DNA replication forks requires the PriA DNA helicase. PriA recognizes and remodels abandoned DNA replication forks performing the DNA unwinding in 3’ to 5’-direction and facilitates loading of the DnaB helicase onto the DNA to restart replication. The single stranded DNA binding protein (SSB) is typically present at the abandoned forks, but there is gap in the knowledge on the interaction between SSB and PriA protein. Here, we used atomic force microscopy (AFM) to visualize the interaction of PriA with DNA substrates in the absence or presence of SSB. Results show that in the absence of SSB, PriA binds preferentially to a fork substrate with a gap in the leading strand. Preferential binding occurs only on forked DNA structures as 5’- and 3’-tailed duplexes were bound equally well. Furthermore, in the absence of SSB, PriA bound exclusively to the fork regions of substrates. In contrast, fork bound SSB loads PriA onto the duplex DNA arms of forks. When the fork has a gap in the leading strand, PriA localizes to both the parental and lagging strand arms. When the gap is present in the lagging strand, PriA is loaded preferentially onto the leading strand arm of the fork. Remodeling of PriA requires a functional C-terminal domain of SSB.

---

A stable DNA replication system is essential for cell viability because DNA replication frequently encounters damages or replication blocks that need to be repaired or removed as soon as possible (1-5). In bacteria, the restart of the replication machinery is mediated by a series of enzymes including the PriA helicase (6,7). It is known that PriA is required for replication restart and that the helicase activity is not required for restart (8). The strong dependence on PriA indicates that a replisome needs to be reassembled at the inactivated forks by a mechanism that can be distinguished from the original initiation by DnaA at *oriC* (9).

PriA is a primosome assembly protein with 3’ to 5’ DNA helicase activity which was discovered because of its requirement for the synthesis of the complementary strand of bacteriophage φX174 single-stranded DNA *in vitro* (10-13). PriA has a two-domain architecture: an N-terminal DNA binding domain (DBD) and a C-terminal helicase domain (HD) (14-18). The cooperation among each domain preserves the recognition and binding activity of PriA to various DNA constructs as well as the interaction with other proteins (6,19,20).

In addition to structure-specific DNA binding, PriA interacts with the single-stranded DNA binding protein (21-23). The single-stranded DNA binding protein (SSB), an essential protein in *E. coli*, binds to and hereby stabilizes the ssDNA that occurs in DNA replication, recombination and repair (24,25). PriA recognizes abandoned DNA replication forks with either duplex DNA or SSB-coated ssDNA and processes these to expose ssDNA as a binding site for primosome components (PriB and DnaT), followed by the reloading of the replicative helicase DnaB (20,26). It was also shown that the PriA helicase activity could be stimulated by binding of SSB onto the initial DNA substrate (14,21). Recent work has shown that the SSB-interaction with PriA involves the oligosaccharide-oligonucleotide binding fold (OB-fold) within the N-terminal domain of the helicase and the linker domain of SSB (27). The mechanism of binding is identical to that between the RecG OB-fold and the linker domain of SSB (28-30).

To understand how PriA interacts with forks and how binding might be influenced by SSB, we used atomic force microscopy (AFM). The experiments were performed in the absence of ATP in order to separate the PriA-DNA binding properties of the protein from its helicase activity. AFM studies revealed the role of the fork type on the efficiency of PriA binding. Furthermore, SSB protein interacts with PriA changing the protein conformation allowing for binding of PriA to the DNA duplex. Experiments with SSB mutant revealed the role of C-terminal segment in this remodeling activity of SSB. These findings are consistent with the ability of SSB protein to remodel RecG that we discovered and described previously (31,32).

## Results

### Binding preference of PriA to various DNA constructs

#### DNA constructs

Fig. 1 shows a set of four DNA constructs used in this work. Constructs T3 and T5 are DNA duplexes with ssDNA tails, with different polarities on different constructs. The use of these two substrates will allow us to elucidate the effect of ssDNA polarity in the assembly of complexes with PriA. The T3 DNA substrate consists of a 244 bp duplex region and a 3’-end 69-nucleotide single-stranded region. The T5 DNA substrate has a 5’-ssDNA region of the same length, but the duplex size is 376 bp. The fork substrates F3 and F5 also differ in the polarity of 69-nt ssDNA. The interaction of PriA with these substrates will allow us to evaluate the role of fork orientation at the junction on the assembled complexes with PriA. F3 construct has a 673-bp duplex region, with a gap in the nascent leading strand. The duplex length of F5 is 675 bp, with a gap in the nascent lagging strand. These forks contained a 69 nt ssDNA arm asymmetrically placed within the DNA duplex region. Consequently, for F3, the DNA duplex regions are a 280 bp parental-strand duplex and a 393 bp lagging-strand duplex. While in F5, the parental-strand duplex has 416 bp residues and the other duplex region is a 260 bp leading-strand duplex. Assembly of each DNA substrate was verified by contour length measurements (Fig. S1).

**Figure 1.**
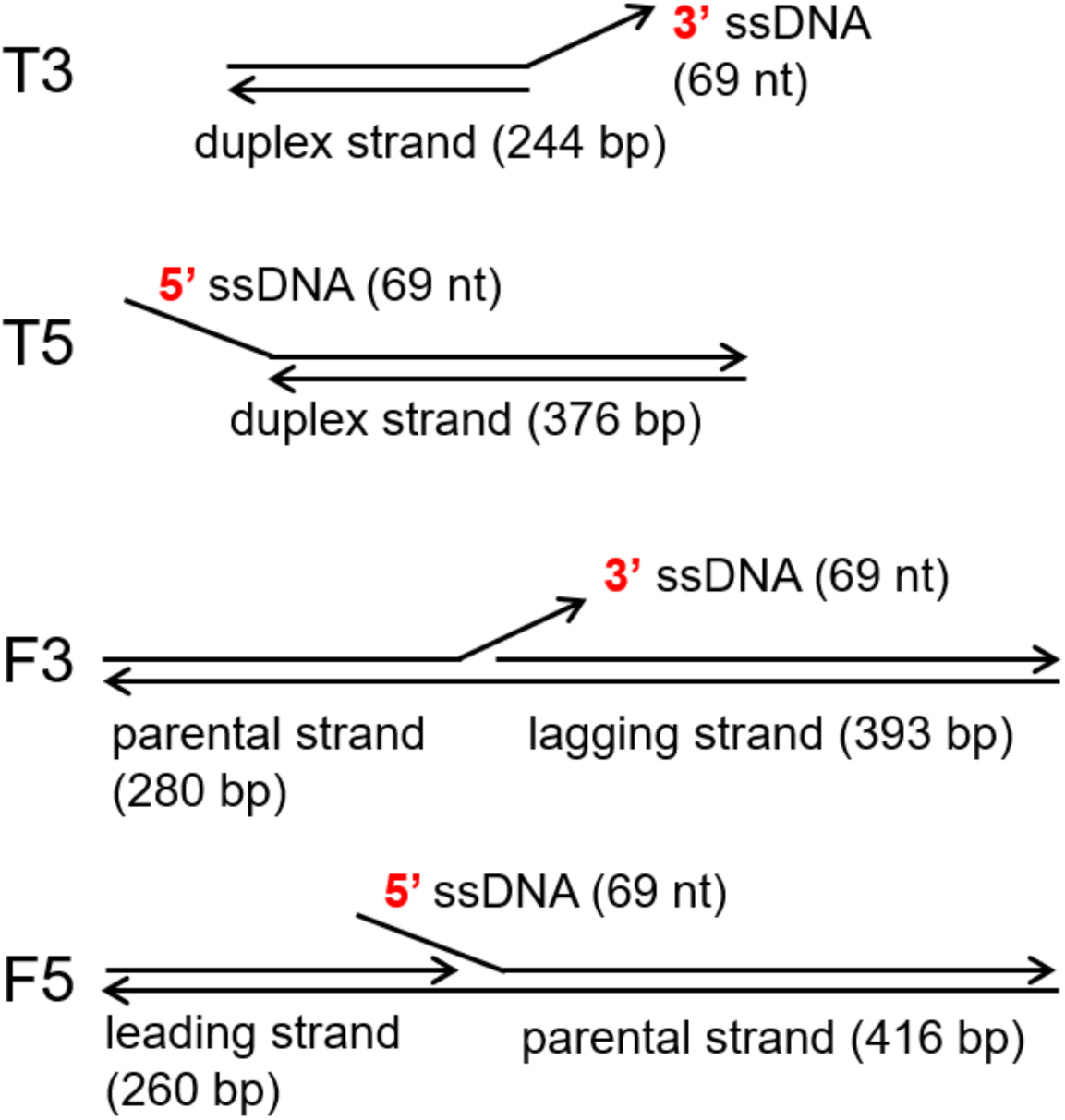
DNA constructs used in this work. T3 DNA construct comprises a 244 bp duplex region and a 3’-end 69-nucleotide single-stranded region. T5 DNA construct has a 5’-ssDNA region of the same length, but the duplex size is 376 bp. In F3 (3’-end) and F5 (5’-end) DNA, 69 nt ssDNA was placed inside the 673 bp (676 bp in F5) duplex with unequal lengths of the DNA duplex regions. Arrows mark the 3’ ends of DNA strands.

#### Interactions of SSB protein with the DNA substrates

To characterize the substrates and evaluate the accessibility of ssDNA, SSB protein was separately bound to each of the substrates. AFM images are shown in Fig. S2A and S2B, which demonstrate that SSB binds to only one of the two ends on tail DNA substrates. On fork DNA substrates, the position of SSB was measured from the end of the shorter arm towards the center of the blob in each DNA-SSB complex (Fig. S2E). The data shown as histograms in this figure were approximated by single-peak Gaussians. The peak values, 276 ± 14 bp (SD) on F3 DNA and 260 ± 16 bp on F5 DNA, correspond to the fork position which are 280 bp and 260 bp for F3 and F5 constructs, respectively. Therefore, and as expected, SSB binds only to the ssDNA regions of each substrate.

#### PriA binding to the tailed DNA substrates

When mixed with T3 and T5 DNA substrates, PriA binds to only one end of the tail DNA substrates regardless of the large excess of the protein (8:1 PriA-to-DNA molar ratio; Fig. 2A and 2B, large-scale images shown in Fig. S3A and S3B). Analysis of over 500 PriA-DNA complexes demonstrated that in <0.5% PriA was observed bound to the both the ssDNA tail and the blunt end. This suggests that PriA binds poorly to either blunt ends or dsDNA. Furthermore, the binding yield of PriA on tail DNA substrates, collected from three independent experiments, was 9.2 ± 0.3% on T3 DNA and 7.9 ± 0.7% on T5 DNA, pointing to a minor preference for PriA binding to the substrate with 3’-ssDNA tail.

**Figure 2.**
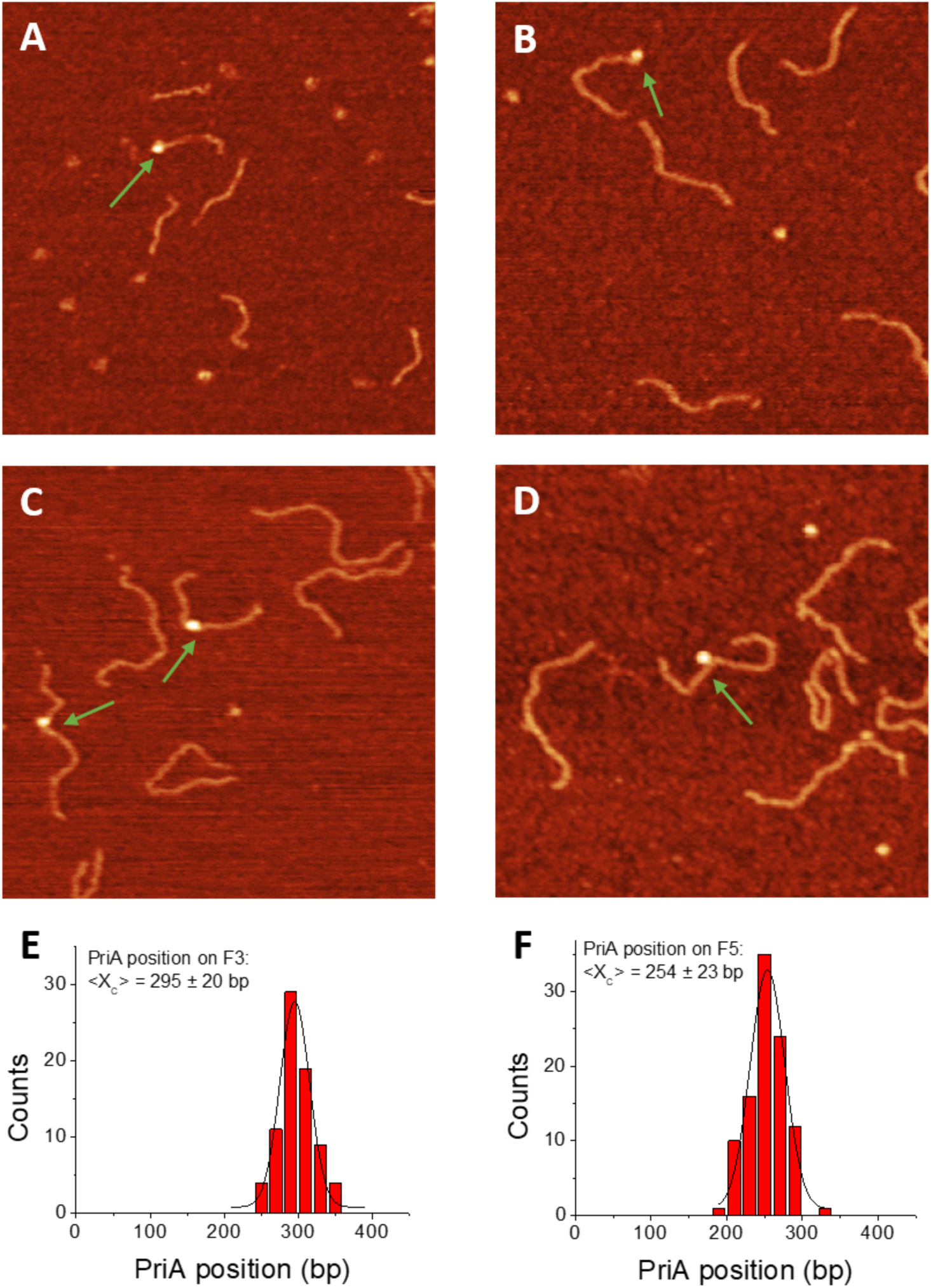
In the absence of SSB, PriA binds preferentially to F3. (A) – (D) 0.5 × 0.5 µm AFM images of T3 with PriA, T5 with PriA, F3 with PriA, and F5 with PriA, respectively. Arrows direct to bound PriA on DNA substrates. Z-scale is 3 nm. (E) and (F) Histograms for PriA position on fork DNA substrates, approximated by Gaussian with bin size of 20 bp. The peaks were found to be centered at 288 ± 24 bp (S.D.) on F3 DNA and at 254 ± 23 bp on F5 DNA, respectively.

#### PriA binding to the fork DNA substrates

To assess the interactions of PriA with fork DNA substrates, PriA was separately bound to F3 and F5 fork DNAs and imaged (Fig. 2C and D). The results show that protein was bound exclusively to a site embedded within the duplex region. To determine if the binding site corresponds to the position of the fork, we measured the position of PriA in each complex, from the end of the shorter arm toward the center of the blob (protein). The histograms, shown in Fig. 2E and 2F, were fitted by single-peak Gaussians with bin size of 20 bp. The centered peak of histogram for PriA position on F3 was found to be 295 ± 20 bp (S.D.). For F5-PriA complexes, the peak was centered at 254 ± 23 bp (S.D.). These peak values match the designed fork position, which support our previous assumption that the fork provides the recognition and binding site for PriA to load onto DNA replication forks.

The yields of complexes for both substrates were measured, and the data revealed different results compared with tailed DNA substrates. The yield of PriA on F3 DNA was 13.0 ± 1.2% while it was 8.2 ± 1.3% on F5 DNA based on the results of three independent experiments. As the polarity of ssDNA tail does not play an essential role in the binding preference for PriA, the difference between F3 and F5 DNA suggests that other structural features of the fork substrates must be involved.

### Interactions between the SSB protein and PriA on fork DNA substrates

Studies show that SSB binds to the RecG and PriA helicases both *in vivo* and *in vitro* (21-23). Previously, we demonstrated that SSB remodeled RecG during DNA loading (31,32). To determine if SSB modulates PriA loading, a similar analysis was performed. Here, PriA was preincubated with SSB at a 2:1 molar ratio for 10 minutes on ice. The mixture was added to fork DNA substrates in the molar ratio of 2:1 (complex: DNA substrate), allowed to bind for 10 minutes at room temperature and imaged.

#### SSB loading PriA onto duplex regions of fork DNA substrates

The resulting images show that for both F3 and F5, double-blob complexes were observed (Fig. 3A and B). The double-blobs correspond to PriA and SSB bound to the same DNA molecule, with the larger blob being SSB and the smaller, PriA, as explained below. Furthermore, in some complexes, the two blobs are located far from each other, while in others, the two blobs co-localize on the DNA (Fig. 3, enlarged images). These data suggest that interaction of PriA with SSB leads to the change PriA conformation allowing for the protein to bind to DNA duplex. We termed this property of SSB remodeling, which was initially identified for SSB mediated loading of RecG protein on the DNA fork.

**Figure 3.**
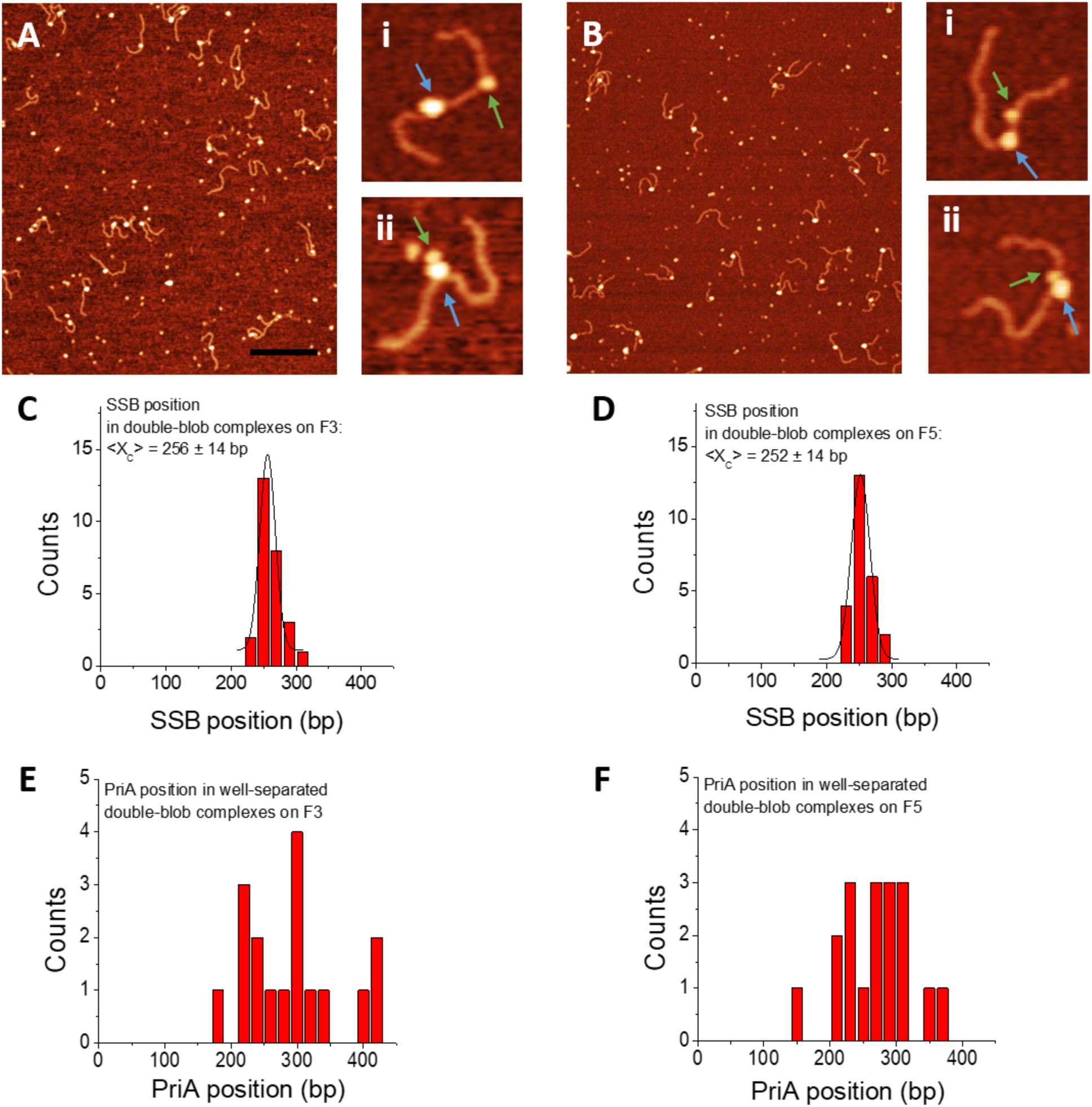
In the presence of SSB, PriA can be localized to duplex regions of forks. (A) and (B) Large-scale AFM images of F3+SSB+PriA and F5+SSB+PriA, respectively. Bar size is 300 nm. Z-scale is 3 nm. (i) and (ii) Gallery of zoomed-in images of each complex to the right. Green arrows direct to PriA in the complexes, while SSB position is directed by blue arrows. (C) and (D) Histograms for SSB position in double-blob complexes, fitted by Gaussian function with bin size of 20 bp. Peaks of SSB distributions were approximated at 256 ± 14 bp (S.D.) on F3 DNA and at 252 ± 14 bp on F5 DNA, respectively. (E) and (F) Histograms for PriA position in well-separated double-blob complexes on F3 DNA and F5 DNA, respectively.

To determine the identity of each blob, PriA and SSB were bound to each DNA substrate separately, imaged, and blob sizes measured. As measured, SSB volume is between 159 ± 29 and 168 ± 27 nm^3^ (Fig. S2). In contrast, the volume values for PriA range from 113 ± 23 to 116 ± 29 nm^3^ (Fig. S3). Therefore, and as observed previously for RecG, the large blobs correspond to SSB and the smaller ones to the DNA helicase. Using this information, the position of binding of SSB and of PriA could be determined using the same measurement approach used for PriA only. The results show that for forks F3 and F5, SSB is bound at the fork (Fig. 3C ad D). The distribution of SSB positions is narrow and the Gaussian maximum (F3: 256 ± 14 bp and F5: 252 ± 14 bp) correlates with specific binding of SSB to the ssDNA region at the fork.

To ascertain PriA binding positions on both flanks of the fork substrates, the locations of the smaller blobs were mapped relative to that of SSB. The results show that in contrast to PriA alone, SSB enables loading of PriA onto the duplex regions flanking the fork as well as at the fork (Fig. 3E and 3F). A summary of positions is presented in Fig. 4. Here, SSB position (green squares) is set to zero, marking the fork. For fork F3, PriA (red dots) is observed bound to the negative interval corresponding to the parental duplex arm, and positive values indicate PriA positioning on the lagging strand. The occurrence of PriA positioning was found to be 28, 32 and 40%, corresponding to parental arm, lagging strand, or co-localizing with SSB at the fork, respectively. Therefore, in the presence of SSB, PriA is loaded preferentially at the fork, and while it is loaded onto the duplex regions as well, there is no clear preference for one region over the other. In contrast, for fork F5, the occurrence of PriA positioning was found to be 24, 48 and 28%, corresponding to parental, leading strand, or co-localizing with SSB, respectively. Therefore, when the fork has a gap in the nascent lagging strand, PriA is preferentially loaded onto the leading strand of the fork.

**Figure 4.**
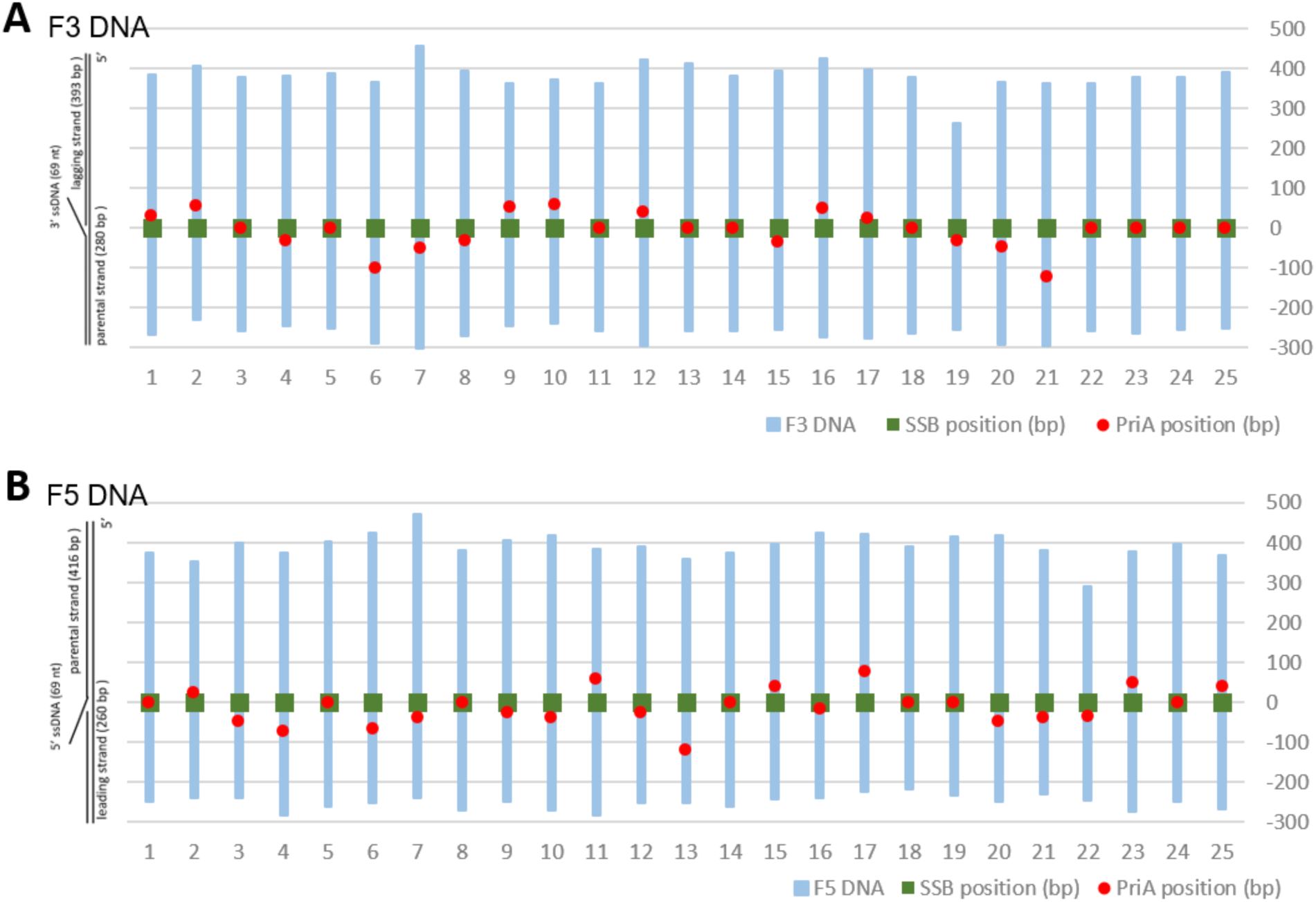
The distributions of proteins in double-blob complexes on each fork DNA substrate, with the SSB position corresponding to zero value on the maps. Green squares indicate the position of SSB, and the red dots point to PriA position. (A) Map of proteins on F3 DNA substrate. PriA in negative interval means that they positioned on the parental strand, and positive values indicate PriA positioning on the lagging strand. (B) Map of proteins on F5 DNA substrate. PriA in negative interval means that they positioned on the leading strand, and positive values indicate PriA positioning on the parental strand.

#### SSB-PriA assembly at the fork

At the fork position, in addition to the colocalized two-blob complexes, single-blob complexes were also observed but the sizes of these single blobs varied. The single blob could be PriA or SSB only, or SSB-PriA complexes, which are of larger sizes. Volume analysis by cross-section was performed to identify the possible component of each single-blob complex. The volume distribution fitted with multi-peak Gaussian for single-blob complexes in F3 DNA sample is shown in Fig. 5A. Peak 1 is centered at 137 ± 37 nm^3^, which is close to bound SSB volume on F3 DNA (X_c_ = 159 ± 29 nm^3^ in Fig. S2H). Peak 2 is approximated at 244 ± 28 nm^3^, corresponding to the complexes of PriA (free protein volume 58 ± 12 nm^3^, shown in Fig. S5A) and SSB (SSB volume bound to DNA is 159 ± 29 nm^3^). The population of those large blobs is ∼24% approximated by the area under Gaussian.

**Figure 5.**
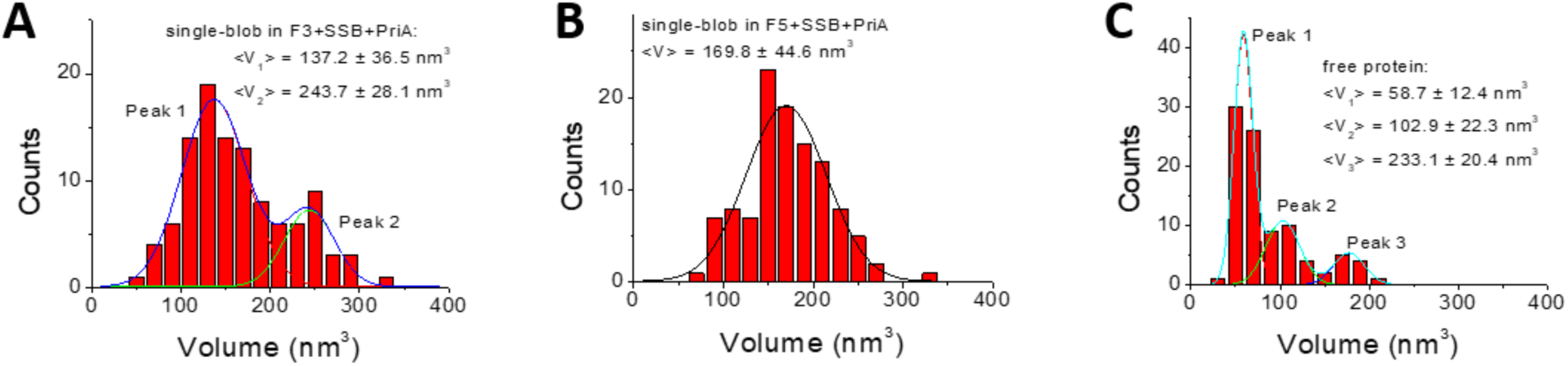
Volume analysis for samples of fork DNA mixed with SSB and PriA. (A) Multi-peak Gaussian fitted distribution for single-blobs on F3 DNA. Peak 1 is centered at 137.2 ± 36.5 nm^3^ and Peak 2 is approximated at 243.7 ± 28.1 nm^3^. The population approximated by area under curve is 76.1% for Peak 1 and 23.9% for Peak 2. (B)The volume distribution for single-blobs on F5 DNA. It was fitted with single-peak Gaussian with a centered peak at 169.8 ± 44.6 nm^3^. (C) Volume distribution for free proteins in samples of fork DNA mixed with SSB and PriA. The histogram shows three peaks based on multi-peak Gaussian fitting, which are 58.7 ± 12.4 nm^3^, 102.9 ± 22.3 nm^3^ and 233.1 ± 20.4 nm^3^. The population approximated by area under curve is 59.9% for Peak 1, 27.5% for Peak 2 and 12.6% for Peak 3.

Similar analysis was done for F5 substrates with data shown in Fig. 5B. The volume distribution was fitted with single-peak Gaussian with a centered peak at 170 ± 45 nm^3^. This peak value shows minor difference from the volume of bound SSB on F5 (168 ± 27 nm^3^, shown in Fig. S2I), which indicates that the interaction of SSB and PriA at the fork region of F5 is subtle.

### The role of the C-terminal domain of SSB in PriA loading

It is known that the C-terminal domain of SSB is required for partner binding (21,28). To determine whether the C-terminal domain of SSB is required for PriA loading, we used the SSBΔC8 protein which has the acidic tip removed. First, the binding of the SSB mutant to forks F3 and F5 was assessed. Results show that yield of SSBΔC8 binding onto each DNA substrate was 83.7% for F3 DNA and 81.8% for F5 DNA (Fig. S4). This is within experimental error, the same as that observed for wild type. Thus, as observed previously, ssDNA binding for this mutant is unaffected (27,33,34).

In mixing experiments, PriA and SSBΔC8 were mixed as for wild type, bound to the DNA and imaged. The yield of double-blob complexes formed on F3 and F5 was reduced to less than 5%, which is lower than that of WT SSB and PriA (table 1). Furthermore, analysis of the sizes of the single blob complexes revealed that these contained SSBΔC8 only as the sizes 97 ± 26 and 111 ± 31 nm^3^ on F3 and F5, respectively (Fig. 6). Collectively these results show that SSBΔC8 does not facilitate loading of PriA onto the DNA.

**Table 1.** Binding yield of DNA with PriA at the molar ratio of 1:8.

|  | **T3+PriA** | **T5+PriA** | **F3+PriA** | **F5+PriA** |
| --- | --- | --- | --- | --- |
| **Binding yield** | **9.2 ± 0.3%** | **7.9 ± 0.7%** | **13.0 ± 1.2%** | **8.2 ± 1.3%** |

**Figure 6.**
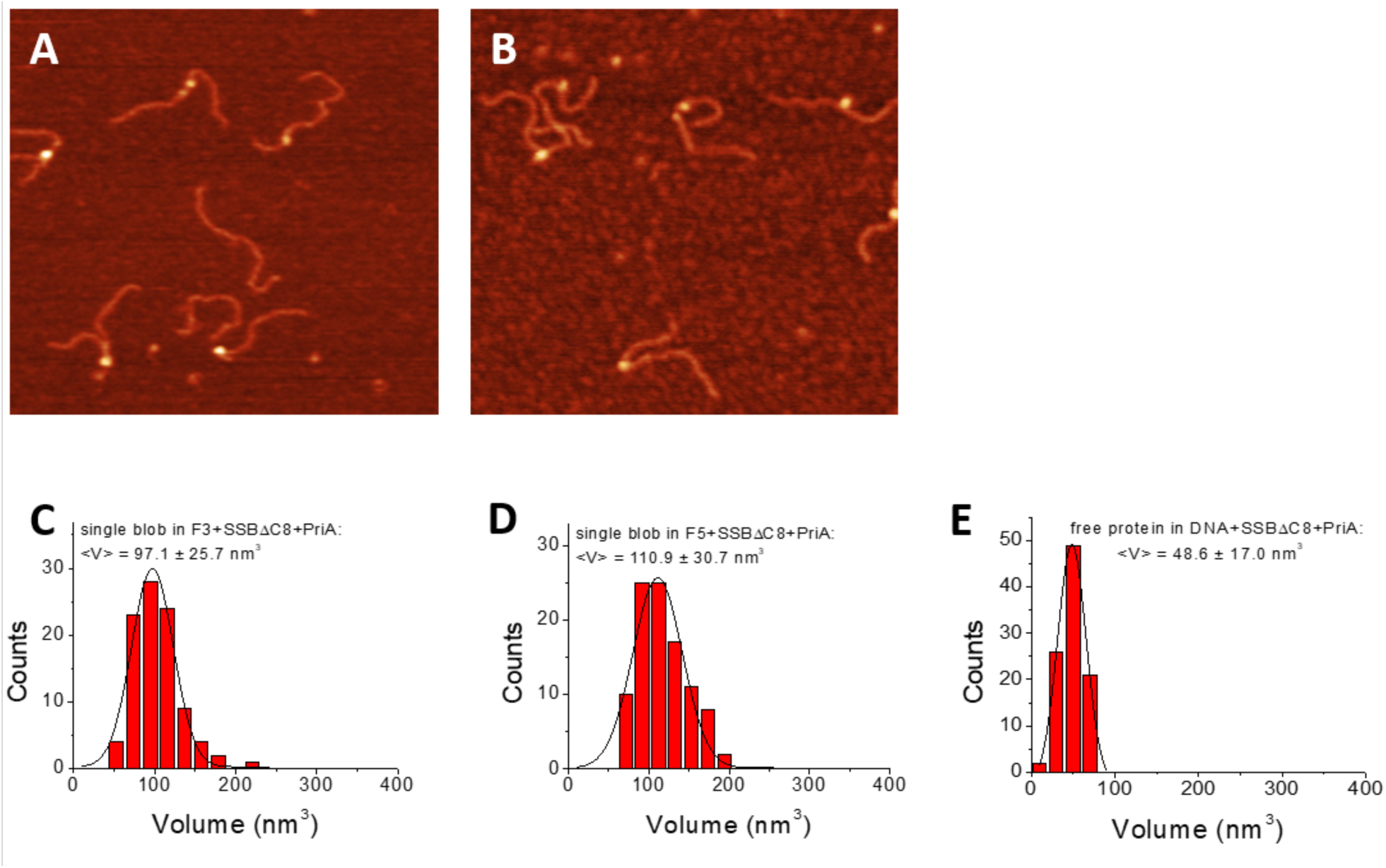
SSBΔC8 does not load PriA. (A) and (B) 0.5 × 0.5 µm AFM images of F3 with SSBΔC8 and PriA, F5 with SSBΔC8 and PriA, respectively. Z-scale is 3 nm. (C) Volume analysis of single-blobs on F3 DNA. The distribution was fitted with single-peak Gaussian and the peak was found to be centered at 97.1 ± 25.7 nm^3^. (D) Single-peak Gaussian fitted volume distribution for single-blobs on F5 DNA. The centered peak is approximated at 110.9 ± 30.7 nm^3^. (E) The volume distribution for free proteins in the sample of fork DNA substrates mixed with SSBΔC8 mutant and PriA. The histogram was fitted by Gaussian with a single peak centered at 48.6 ± 17 nm^3^.

We also characterized the role of C-terminus on interaction of SSB and PriA in the absence of DNA. The analysis was made for free proteins present in the same samples on the protein-DNA sample characterized above. Size analysis for free proteins in the sample of fork DNA substrates mixed with SSBΔC8 mutant and PriA is shown in Fig. 6E. The volume distribution was fitted by Gaussian with a single peak centered at 49 ± 17 nm^3^, which is different from multi-peak distribution of free proteins in samples of fork DNA substrates with WT SSB and PriA mixture. Thus, SSB and PriA interaction was observed in the absence of DNA by size analysis, while the difference between results of PriA with WT SSB and SSBΔC8 mutant further emphasizes the role of SSB C-terminus in the interactions with its partner protein. Thus, C-terminus of SSB is needed for SSB-PriA interaction.

### Interaction between SSB and PriA in the absence of DNA

To characterize the assembly of SSB-PriA complexes in the absence of DNA, we also analyzed sizes of free proteins in the samples of fork DNA substrates mixed with SSB and PriA. The volume distribution is shown in Fig. 5C. The histogram shows three peaks based on multi-peak Gaussian fitting, which are 59 ± 12 nm^3^, 103 ± 22 nm^3^ and 233 ± 20 nm^3^. Peak 1 matches the volume of free PriA (58 ± 12 nm^3^), shown in Fig. S5A. Peak 2 is close to the volume of free SSB (91 ± 21 nm^3^ in Fig. S5B). Peak 3 could be assigned to the SSB-PriA complex even though the measured volume of each protein was smaller than half of the 3^rd^ peak value. Because neither tetrameric SSB nor monomeric PriA aggregates with themselves, based on the narrow single-peak distribution of each volume measurement in our control experiments (shown in Fig. S5A and B). Note that Peak 3 is with a smallest probability, which is 12.6% evaluated by the area under Gaussian. While the probabilities of the other two are 59.9% for Peak 1 and 27.5% for Peak 2. Therefore, though the population of the SSB-PriA complexes is low, the interaction between SSB and PriA in the absence of DNA also exists.

## Discussion

The major function of PriA, a DNA helicase, is to restart the replication process by facilitating the loading of DNA polymerase onto the stalled replication fork. Once bound, PriA exposes ssDNA for the replicative DNA helicase DnaB by unwinding the lagging strand or by remodeling SSB-coated ssDNA (20,35). This priming process requires the binding of PriA to various DNA structures (6), which was the focus of this paper, as it remains unclear how PriA differentiates among various DNA structures and then plays the role that is needed for the restart of stalled replication fork is still not well studied. To uncouple the DNA binding property of PriA from its helicase activity we performed studies in the absence of ATP. The major findings are discussed below.

### The effects of ssDNA polarity and the fork structure on the interaction between PriA and DNA

On the fork DNA substrates, PriA binds to both substrates at the fork position, with a preference to the fork DNA substrate with a gap in the nascent leading strand (F3), which is an unexpected finding. According to other publications (17,20,35,36), PriA should bind preferentially to fork F5 (has a gap in the nascent lagging strand), since F5 has a 3’-OH group at the fork position as the recognition site for the 3’ binding domain in PriA, while F3 does not. Our finding on PriA’s preference for binding suggests the participation of the other domains in PriA-DNA binding, and is in line with the previous finding that PriA binds to the arrested replication fork in a manner independent on 3’-terminus as well (17). In addition to the independence of polarity, the bend at the fork position may play an important role in PriA binding activity as well, as demonstrated in refs. (14,37). This emphasizes the essential role of the fork structure and the presence of a nascent lagging strand in the binding of PriA onto stalled replication fork.

Our model on the role of other PriA domains in assembly of the complexes with the replication fork is supported by the findings that PriA does not show binding preference to the ssDNA of tail DNA substrates (Fig. 2A, B; Table 1). P Nurse et al. (37) demonstrated that PriA bound with high affinity to duplexes with 3’-tails, whereas it did not bind to duplexes with 5’-tails at all, however, they detected stable binding of the 3’-extension when the ssDNA tail exceeded 12 nt and high affinity binding resulted when the tails were in excess of 16 nt in length. In our studies the size of ssDNA region is 69 nt. The excess of ssDNA region might also involve in binding activity of multidomain and then stabilize the interaction between PriA and DNA substrates. Note as well that PriA has over 100-time higher binding yield with the ssDNA region compared to the blunt end on the tail DNA substrates. This property also contributes to the high efficiency of targeting at the stalled replication fork where the restart is needed.

### Remodeling of PriA by SSB on fork DNA substrates

Although interactions between SSB protein and PriA has already been characterized (14,21,22,38,39), our results reveled a novel role of SSB in interaction of PriA with fork DNA substrates. Shown in Fig. 3A and B, both proteins can be co-localized with the fork position (insets (ii)) in the AFM images, which has PriA bound to the fork while SSB bound to the ssDNA region of the fork. Interestingly, we also identified complexes shown as insets (i) on both images in which SSB and PriA are well separated. On both substrates, SSB locations produce narrow distributions (histograms C and D in Fig. 3). However, positions of PriA in the SSB-PriA complexes assembled on both substrates are very broad (Fig. 3E and D). These data are in contrast with the data obtained from PriA-DNA complexes in the absence of SSB, which has narrow distribution on each histogram and the peak positions coincide with the location of the fork (Fig. 2E and F). We hypothesize that after remodeling, PriA is capable of binding to DNA duplex with spontaneous translocation over DNA duplexes.

The mapping of positions of proteins on the fork substrates in Fig. 4, shows that there is no clear preference of the PriA position on both flanks of the forks for F3 forks. However, on fork F5, which has a gap in the lagging strand, remodeled PriA showed a preference of binding to the leading strand. These suggest that interaction of PriA with SSB changes the helicase conformation in such a way that PriA becomes capable of binding to DNA duplex. Note that in the absence of SSB, such change in DNA binding conformation of PriA has not been identified. Similar remodeling property of SSB effect was found for RecG protein by us (32), and the translocation mobility of RecG has been proven by direct visualization of RecG mobility using time-lapse high-speed AFM approach (31). It was hypothesized that remodeling of RecG by SSB allows RecG to translocate along the duplex strands in an ATP independent way, so that RecG can be recruited rapidly to accomplish its fork regression role (31,32). Recently the protein-protein interaction of SSB with partner protein has been reported for RecQ (40-42) and RecOR (43,44), suggesting that the remodeling of components of the DNA replication machinery is a common property of SSB. Remodeling of PriA by SSB in the absence of ATP increases the ability of PriA binding onto duplex DNA, which was detected as the isolated CTD activity (20). Thereby, the binding and/or translocation of PriA along the duplex may stimulate the loading of PriA onto the stalled replication fork in an ATP-independent way, facilitating the restart process once the ATP is available for PriA helicase activity.

Studies showed that most of the interactions between SSB and its partner protein are mediated by the C-terminus of SSB (22,45). The C-terminal domain of SSB, corresponding to residues 117-178, can be sub-divided into the intrinsically disordered linker (aa 117-170) and the highly-conserved acidic tip (30). The linker mediates protein-protein interactions while the acidic tip is required to maintain the structure of the C-terminus of SSB so that is does not bind to SSB itself (27). Thus, when the tip is mutated or deleted, the C-terminus binds to SSB, thereby inactivating the protein. Consequently, SSB-partner interactions are lost. Experiments with the SSBΔC8 in which the acidic tip of SSB was removed showed that the yield for double-blob complexes dropped and these findings are in line with the effect C-terminus on remodeling of RecG (32), suggesting that C-terminus plays an important role of the SSB remodeling property.

### Protein-protein interactions in the colocalized SSB-PriA complexes

PriA interacts with SSB both in the absence and presence of fork DNA substrates. According to published data, the stimulation effect/localization activity of SSB-PriA interaction requires the excess of SSB over the helicase (14,21,22,38). In our experiments (Fig.s 3A and B), colocalized PriA-SSB complexes on DNA were detected at the molar SSB-to-PriA ratio of 1:2. Importantly, the concentration of PriA was 5 nM. The volume analysis of colocalized SSB and PriA complexes (Fig. 5) was clearly seen for F3 substrate from Peak 2 and the yield of such complexes was 24% compared with the single blob complexes (76%), which are corresponding to complexes with SSB binding only. Considering that the single-blob complexes count for 78% binding events in the mixture sample (with the rest of events being 7% of free DNA substrates and 15% of double-blob complexes), the overall binding yield of PriA-SSB on F3 DNA should be 24% * 78%, which is 18.7%. No such clearly identified peak appears for F5 substrate (Fig. 5B), suggesting that the colocalization depends on the substrate type and is clearly higher for F3 substrate which is a better substrate for binding PriA only (Table 1). In this way, the interaction between SSB and PriA increased the binding of PriA to fork DNA substrates and improved the selectivity of PriA to a more favorable substrate. This suggests that SSB stimulation effect, as it is known on PriA helicase activity, also plays a role in the binding of PriA onto DNA substrates.

In addition to the protein-protein interaction found in protein-DNA complexes, the volume analysis of free protein (Fig. 5C) revealed a minor peak (Peak 3), which has a larger volume than each protein itself, corresponding to the volume of the complex formed from SSB-PriA assembly. So, SSB-PriA interaction is independent of the existence of DNA or ATP. This property helps SSB locate or direct PriA to the place it is needed in a more efficient way.

Conclusions: Our AFM studies revealed several novel properties of interactions between PriA and stalled DNA replication forks, with or without SSB. In the absence of ATP, we observed that PriA binds preferentially to the forked DNA with a gap in the nascent leading strand, compared to the other forked DNA substrate which has a gap in the nascent lagging strand. Since PriA showed no clear preference towards the polarity of ssDNA in tail DNA substrates, it is the fork structure that plays a more essential role in PriA binding. The interactions between SSB and PriA revealed the remodeling of PriA by SSB, which loaded PriA onto the duplex DNA, and this property of SSB can be attributed to its C-terminal segment. It could be an ATP-independent translocation activity by which PriA slides along the DNA duplex and search for the site needed to get restarted.

## Experimental procedures

### Protein preparation

Purification of the PriA protein followed the method described previously (22). The *his-PriA protein* was purified by ammonium sulfate precipitation followed by affinity chromatography using HisTrap FF crude column, SP Sepharose column (Equilibrated with 20 m*M* potassium phosphate, pH 7.6, 150 m*M* KCl, 0.1 m*M* EDTA, and 1 m*M* DTT; Eluted with a linear 150–500 m*M* KCl gradient) and Heparin column (Equilibrated with 20 m*M* Tris-OAc, pH 7.5, 0.1 m*M* EDTA, 1 m*M* DTT, 10% (v/v) Glycerol, and 100 m*M* KCl; Eluted with a linear KCl gradient of 100–600 m*M*). Fractions containing PriA were pooled and dialyzed overnight against storage buffer (20 m*M* Tris-HCl (pH 7.5), 1 m*M* DTT, 400 m*M* KCl, and 50% (v/v) glycerol). The PriA concentration was determined using extinction coefficient of 104,850 M^−1^ cm^−1^ (46).

SSB protein was purified from strain K12ΔH1Δtrp, as described in ref (25,47). The concentration of the purified protein was determined at 280 nm using *ε* = 30,000 M^−1^ cm^−1^ (46).

### DNA substrates preparation

Each tail DNA substrate (T3 or T5) was assembled from a duplex-DNA segment and a tail-DNA segment. The fork DNA substrates (F3 and F5) were assembled from two duplex-DNA segments and a core fork segment as described previously (31,32). The two duplex-DNA segments (224 bp segment and 356 bp segment) were precisely the same as the segments we had in ref (31,32). To assemble the tail-DNA segment for T3, the ssDNA oligonucleotides O42 (5’-TCATGACTCGCTGCGCAAGGCTAACAGCATCACACACATTAACAATTCTAACATCTG-3’) and O43 (5’-CCTTGCGCAGCGAGTCA-3’) were phosphorylated, mixed in an equal molar ratio, and then annealed by heating to 95°C in the hot boiled water and then cooling down slowly to room temperature. The tail-DNA segment was then ligated with the 224bp duplex-DNA segment in the molar ratio of 1:1 to assemble the T3 DNA substrate. Similar to the assembly of T3 DNA substrate, the tail-DNA segment for T5 was annealed from the 5’-phosphorylated oligonucleotides O36 (5’-TACGTGTAGGAATTATATTAAAGAGAAAGTGAAACCCAAAGAATGAAAAAGAAGATGTTAGAATTGTAAGCGGTATCAGCTCACTCATA-3’) and O37 (5’ GCTTATGAGTGAGCTGATACCGC-3’) in the equal molar ratio and then was ligated together with the 356 bp duplex-DNA segment in 1:1 molar ratio to obtain the T5 DNA substrate. The core fork segment of F3 was assembled by annealing the 5’-phosphorylated oligos in the same molar ratio (O30: 5’-TCATCTGCGTATTGGGCGCTCTTCCGCTTCCTATCT-3’; O31: 5’-TCGTTCGGCTGCGGCGAGCGGTATCAGCTCACTCATA-3’; O32: 5’-GCTTATGAGTGAGCTGATACCGCTCGCCGCAGCCGAACGACCTTGCGCAGCGAGTCAGTGAGATAGGAAGCGGAAGAGCGCCCAATAC GCAGA-3’ and O33: 5’-CACTGACTCGCTGCGCAAGGCTAACAGCATCACACACATTAACAATTCTAACATCTGGGTTTTCATTCTTTGGGTTTCACTTTCTCCAC-3’). While the core fork segment of F5 was annealed from phosphorylated O30, O31, O32, and O34 (O34: 5’-CTAACAGCATCACACACATTAACAATTCTAACATCTGGGTTTTCATTCTTTGGGTTTCACTTTCTCCACCACTGACTCGCTGCGCAAGG-3’), in the same molar ratio. The two duplexes and core fork segment were ligated together at the molar ratio of 1:1:1 at 16°C overnight. The final products were purified with HPLC using a TSKgel DNA-STAT column. All oligonucleotides were bought from IDT (Integrated DNA Technologies, Inc., Coralville, Iowa, USA).

### Protein-DNA complex and AFM sample preparation

PriA-DNA complex was prepared by mixing the PriA monomer (molar concentration: 100 nM) with DNA substrates (molar concentration: 45 nM) in a molar ratio of 8:1. The mixture was incubated in 10 µl of binding buffer [10mM Tris-HCl (pH 7.5), 50 mM NaCl, 5 mM MgCl_2_, 1 mM DTT] for 10 min at room temperature. After incubation, the complex was then diluted to achieve lower DNA concentration (∼1 nM), which was ready for deposition onto the APS functionalized mica.

SSB-PriA-DNA complex was prepared by mixing the proteins first: the SSB tetramer (molar concentration: 50 nM) was mixed with PriA monomer in a molar ratio of 1:2 and the mixture was kept on ice for 30 minutes before use. The mixture of proteins was added to fork DNA substrates in a 1:2:4 (DNA substrates: SSB: PriA) molar ratio and then incubated in 10 µl binding buffer for 10 min at room temperature. After incubation, the complex was then diluted to achieve lower DNA concentration (∼1 nM) for AFM imaging using APS functionalized mica procedure.

1-(3-aminopropyl) silatrane (APS) functionalized mica was used as the AFM substrate for all experiments (48). Fresh cleaved mica was incubated in 4ml APS (167 µM) in a cuvette for 30 min and then was rinsed with ddH_2_O thoroughly as described before (31,32). Ten microliters of the sample were deposited onto the APS functionalized mica for 2 min. After 2-minutes incubation, the mica was rinsed with ddH_2_O, and then was dried with a gentle Argon-gas flow.

### AFM imaging and data analysis

Images were acquired using tapping mode in the air on a MultiMode 8, Nanoscope V system (Bruker, Santa Barbara, CA) using TESPA probes (320 kHz nominal frequency and a 42 N/m spring constant) from the same vendor. The dry sample AFM images were analyzed using the FemtoScan Online software package (Advanced Technologies Center, Moscow, Russia). The positions of each protein were measured from the end of the shorter arm on the DNA substrates towards the center of the protein. The contour lengths of the DNA were then continuously measured from the center of the protein towards the other end of the DNA substrate. The yield of protein-DNA complexes was calculated from the number of complexes dividing by the total number of DNA molecules. The histograms were approximated with Gaussian distribution and the mean values and errors (SD and SEM) were calculated by using Origin software (OriginLab Corporation, Northampton, MA, USA). The protein height and volume were measured with the cross-section option. The volume was calculated by applying the measured data to the formula: V = 3.14 * H/6 * (0.75 * D_1_ * D_2_ * H^2^), in which D_1_ and D_2_ are the diameters of the protein, which were measured twice, and H is the highest height out of the two measurements of the protein (32,49,50).

## Acknowledgements

The work was supported by the National Institutes of Health grants R01 GM118006 to YLL and R01 GM100156 to PRB and YLL.

## Conflict of interest

There are no conflicts to declare.

## FOOTNOTES

Funding was provided by grants R01 GM118006 to YLL and R01 GM100156 to PRB and YLL.

## The abbreviations used are

SSB: the single stranded DNA binding protein;
AFM: atomic force microscopy;
DBD: DNA binding domain;
HD: helicase domain.

## References

1. Cox, M. M., Goodman, M. F., Kreuzer, K. N., Sherratt, D. J., Sandler, S. J., and Marians, K. J. (2000) The importance of repairing stalled replication forks. Nature 404, 37–41

2. Kogoma, T. (1997) Stable DNA replication: interplay between DNA replication, homologous recombination, and transcription. Microbiol Mol Biol Rev 61, 212–238

3. McGlynn, P., and Lloyd, R. G. (2002) Recombinational repair and restart of damaged replication forks. Nat Rev Mol Cell Biol 3, 859–870

4. Gabbai, C. B., and Marians, K. J. (2010) Recruitment to stalled replication forks of the PriA DNA helicase and replisome-loading activities is essential for survival. DNA Repair (Amst) 9, 202–209

5. Michel, B., Sinha, A. K., and Leach, D. R. F. (2018) Replication Fork Breakage and Restart in Escherichia coli. Microbiol Mol Biol Rev 82

6. Windgassen, T. A., Wessel, S. R., Bhattacharyya, B., and Keck, J. L. (2018) Mechanisms of bacterial DNA replication restart. Nucleic Acids Res 46, 504–519

7. Sandler, S. J., and Marians, K. J. (2000) Role of PriA in replication fork reactivation in Escherichia coli. J Bacteriol 182, 9–13

8. Zavitz, K. H., and Marians, K. J. (1992) ATPase-deficient mutants of the Escherichia coli DNA replication protein PriA are capable of catalyzing the assembly of active primosomes. J Biol Chem 267, 6933–6940

9. Michel, B., Grompone, G., Flores, M. J., and Bidnenko, V. (2004) Multiple pathways process stalled replication forks. Proc Natl Acad Sci U S A 101, 12783–12788

10. Wickner, S., and Hurwitz, J. (1975) Association of phiX174 DNA-dependent ATPase activity with an Escherichia coli protein, replication factor Y, required for in vitro synthesis of phiX174 DNA. Proc Natl Acad Sci U S A 72, 3342–3346

11. Zavitz, K. H., and Marians, K. J. (1991) Dissecting the functional role of PriA protein-catalysed primosome assembly in Escherichia coli DNA replication. Mol Microbiol 5, 2869–2873

12. Masai, H., Asai, T., Kubota, Y., Arai, K., and Kogoma, T. (1994) Escherichia coli PriA protein is essential for inducible and constitutive stable DNA replication. EMBO J 13, 5338–5345

13. Arai, K., Low, R. L., and Kornberg, A. (1981) Movement and site selection for priming by the primosome in phage phi X174 DNA replication. Proc Natl Acad Sci U S A 78, 707–711

14. Chen, H. W., North, S. H., and Nakai, H. (2004) Properties of the PriA helicase domain and its role in binding PriA to specific DNA structures. J Biol Chem 279, 38503–38512

15. Tanaka, T., Mizukoshi, T., Taniyama, C., Kohda, D., Arai, K., and Masai, H. (2002) DNA binding of PriA protein requires cooperation of the N-terminal D-loop/arrested-fork binding and C-terminal helicase domains. J Biol Chem 277, 38062–38071

16. Ouzounis, C. A., and Blencowe, B. J. (1991) Bacterial DNA replication initiation factor priA is related to proteins belonging to the ‘DEAD-box’ family. Nucleic Acids Res 19, 6953

17. Tanaka, T., Mizukoshi, T., Sasaki, K., Kohda, D., and Masai, H. (2007) Escherichia coli PriA protein, two modes of DNA binding and activation of ATP hydrolysis. J Biol Chem 282, 19917–19927

18. Tanaka, T., Taniyama, C., Arai, K., and Masai, H. (2003) ATPase/helicase motif mutants of Escherichia coli PriA protein essential for recombination-dependent DNA replication. Genes Cells 8, 251–261

19. Windgassen, T. A., Leroux, M., Sandler, S. J., and Keck, J. L. (2019) Function of a strand-separation pin element in the PriA DNA replication restart helicase. J Biol Chem 294, 2801–2814

20. Bhattacharyya, B., George, N. P., Thurmes, T. M., Zhou, R., Jani, N., Wessel, S. R., Sandler, S. J., Ha, T., and Keck, J. L. (2014) Structural mechanisms of PriA-mediated DNA replication restart. Proc Natl Acad Sci U S A 111, 1373–1378

21. Cadman, C. J., and McGlynn, P. (2004) PriA helicase and SSB interact physically and functionally. Nucleic Acids Res 32, 6378–6387

22. Yu, C., Tan, H. Y., Choi, M., Stanenas, A. J., Byrd, A. K., K, D. R., Cohan, C. S., and Bianco, P. R. (2016) SSB binds to the RecG and PriA helicases in vivo in the absence of DNA. Genes Cells 21, 163–184

23. Costes, A., Lecointe, F., McGovern, S., Quevillon-Cheruel, S., and Polard, P. (2010) The C-terminal domain of the bacterial SSB protein acts as a DNA maintenance hub at active chromosome replication forks. PLoS Genet 6, e1001238

24. Meyer, R. R., and Laine, P. S. (1990) The single-stranded DNA-binding protein of Escherichia coli. Microbiol Rev 54, 342–380

25. Lohman, T. M., and Ferrari, M. E. (1994) Escherichia coli single-stranded DNA-binding protein: multiple DNA-binding modes and cooperativities. Annu Rev Biochem 63, 527–570

26. Allen, G. C., Jr., and Kornberg, A. (1993) Assembly of the primosome of DNA replication in Escherichia coli. J Biol Chem 268, 19204–19209

27. Ding, W., Tan, H. Y., Wilczek, L. A., Zhang, J. X., Mulkin, J. A., Hsieh, K. R., and Bianco, P. R. (2020) The mechanism of SSB-RecG binding: implications for SSB interactome function. Protein Science, submitted

28. Bianco, P. R., Pottinger, S., Tan, H. Y., Nguyenduc, T., Rex, K., and Varshney, U. (2017) The IDL of E. coli SSB links ssDNA and protein binding by mediating protein-protein interactions. Protein Sci 26, 227–241

29. Bianco, P. R., and Lyubchenko, Y. L. (2017) SSB and the RecG DNA helicase: An intimate association to rescue a stalled replication fork. Protein Sci 26, 638–649

30. Bianco, P. R. (2017) The tale of SSB. Progress in biophysics and molecular biology 127, 111–118

31. Sun, Z., Hashemi, M., Warren, G., Bianco, P. R., and Lyubchenko, Y. L. (2018) Dynamics of the Interaction of RecG Protein with Stalled Replication Forks. Biochemistry 57, 1967–1976

32. Sun, Z., Tan, H. Y., Bianco, P. R., and Lyubchenko, Y. L. (2015) Remodeling of RecG Helicase at the DNA Replication Fork by SSB Protein. Sci Rep 5, 9625

33. Tan, H. Y., Wilczek, L. A., Pottinger, S., Manosas, M., Yu, C., Nguyenduc, T., and Bianco, P. R. (2017) The intrinsically disordered linker of E. coli SSB is critical for the release from single-stranded DNA. Protein Sci 26, 700–717

34. Liu, J., Choi, M., Stanenas, A. G., Byrd, A. K., Raney, K. D., Cohan, C., and Bianco, P. R. (2011) Novel, fluorescent, SSB protein chimeras with broad utility. Protein Sci 20, 1005–1020

35. Jones, J. M., and Nakai, H. (2001) Escherichia coli PriA helicase: fork binding orients the helicase to unwind the lagging strand side of arrested replication forks. J Mol Biol 312, 935–947

36. Sasaki, K., Ose, T., Okamoto, N., Maenaka, K., Tanaka, T., Masai, H., Saito, M., Shirai, T., and Kohda, D. (2007) Structural basis of the 3’-end recognition of a leading strand in stalled replication forks by PriA. EMBO J 26, 2584–2593

37. Nurse, P., Liu, J., and Marians, K. J. (1999) Two modes of PriA binding to DNA. J Biol Chem 274, 25026–25032

38. Kozlov, A. G., Jezewska, M. J., Bujalowski, W., and Lohman, T. M. (2010) Binding specificity of Escherichia coli single-stranded DNA binding protein for the chi subunit of DNA pol III holoenzyme and PriA helicase. Biochemistry 49, 3555–3566

39. Cadman, C. J., Lopper, M., Moon, P. B., Keck, J. L., and McGlynn, P. (2005) PriB stimulates PriA helicase via an interaction with single-stranded DNA. J Biol Chem 280, 39693–39700

40. Shereda, R. D., Bernstein, D. A., and Keck, J. L. (2007) A central role for SSB in Escherichia coli RecQ DNA helicase function. J Biol Chem 282, 19247–19258

41. Mills, M., Harami, G. M., Seol, Y., Gyimesi, M., Martina, M., Kovacs, Z. J., Kovacs, M., and Neuman, K. C. (2017) RecQ helicase triggers a binding mode change in the SSB-DNA complex to efficiently initiate DNA unwinding. Nucleic Acids Res 45, 11878–11890

42. Bagchi, D., Manosas, M., Zhang, W., Manthei, K. A., Hodeib, S., Ducos, B., Keck, J. L., and Croquette, V. (2018) Single molecule kinetics uncover roles for E. coli RecQ DNA helicase domains and interaction with SSB. Nucleic Acids Res 46, 8500–8515

43. Hobbs, M. D., Sakai, A., and Cox, M. M. (2007) SSB protein limits RecOR binding onto single-stranded DNA. J Biol Chem 282, 11058–11067

44. Bell, J. C., Liu, B., and Kowalczykowski, S. C. (2015) Imaging and energetics of single SSB-ssDNA molecules reveal intramolecular condensation and insight into RecOR function. Elife 4, e08646

45. Shereda, R. D., Kozlov, A. G., Lohman, T. M., Cox, M. M., and Keck, J. L. (2008) SSB as an organizer/mobilizer of genome maintenance complexes. Crit Rev Biochem Mol Biol 43, 289–318

46. Tan, H. Y., Wilczek, L. A., Pottinger, S., Manosas, M., Yu, C., Nguyenduc, T., and Bianco, P. R. (2017) The intrinsically disordered linker of E. coli SSB is critical for the release from single-stranded DNA. Protein Science 26, 700–717

47. Lohman, T. M., Green, J. M., and Beyer, R. S. (1986) Large-scale overproduction and rapid purification of the Escherichia coli ssb gene product. Expression of the ssb gene under. lambda. PL control. Biochemistry 25, 21–25

48. Shlyakhtenko, L. S., Gall, A. A., and Lyubchenko, Y. L. (2013) Mica functionalization for imaging of DNA and protein-DNA complexes with atomic force microscopy. Methods Mol Biol 931, 295–312

49. Shlyakhtenko, L. S., Lushnikov, A. Y., Miyagi, A., Li, M., Harris, R. S., and Lyubchenko, Y. L. (2012) Nanoscale structure and dynamics of ABOBEC3G complexes with single-stranded DNA. Biochemistry 51, 6432–6440

50. Shlyakhtenko, L. S., Lushnikov, A. Y., Miyagi, A., and Lyubchenko, Y. L. (2012) Specificity of binding of single-stranded DNA-binding protein to its target. Biochemistry 51, 1500–1509

